# Physiological significance of proteolytic processing of Reelin revealed by cleavage-resistant Reelin knock-in mice

**DOI:** 10.1101/2020.01.15.903237

**Authors:** Eisuke Okugawa, Himari Ogino, Tomofumi Shigenobu, Yuko Yamakage, Hitomi Tsuiji, Hisashi Oishi, Takao Kohno, Mitsuharu Hattori

## Abstract

Reelin is a secreted protein that plays versatile roles in neuronal development and function, and hypoactivity of Reelin is implicated in many neuropsychiatric disorders. The strength of Reelin signaling is regulated by proteolytic processing, but its importance *in vivo* is not yet fully understood. Here, we generated Reelin knock-in (PA-DV KI) mice in which the key cleavage site of Reelin was abolished by mutation. As expected, the cleavage of Reelin was severely abrogated in the cerebral cortex and hippocampus of PA-DV KI mice. The amount of Dab1, whose degradation is induced by Reelin signaling, decreased in these tissues, indicating that the signaling strength of Reelin was augmented. The brains of PA-DV KI mice were largely structurally normal, but unexpectedly, the hippocampal layer was disturbed. This phenotype was ameliorated in hemizygote PA-DV KI mice, indicating that excess Reelin signaling is detrimental to hippocampal layer formation. The neuronal dendrites of PA-DV KI mice had more branches and were elongated compared to wild-type mice. These results present the first direct evidence of the physiological importance of Reelin cleavage and suggest that inhibition of Reelin cleavage would counteract neuropsychiatric disorders without causing severe systemic side effects.

## Introduction

Reelin is a large secreted glycoprotein that regulates many important events in mammalian brain development^1–3^. It is also involved in synaptic plasticity and the modulation of higher brain functions^1,4–8^. At the cellular level, Reelin induces the clustering of apolipoprotein E receptor 2 (ApoER2) and very-low-density lipoprotein receptor (VLDLR), which is the prerequisite for the phosphorylation of the cytosolic adaptor protein Dab1^9–11^. Phosphorylated Dab1 is quickly degraded^1,2,5^. Thus, the amount of Dab1 is indicative of the strength of Reelin signaling^12–15^. Several pathways act downstream of Dab1 phosphorylation and regulate the cytoskeleton, adhesion molecules, ion channels, and neurotransmitter release^1–3,5,16^. Accordingly, insufficient Reelin signaling could contribute to the pathogenesis of neuropsychiatric disorders, such as epilepsy^17–19^, schizophrenia^6,16,20^, and autism^3,8,16^. Furthermore, recent studies showed that a decrease or insufficiency in Reelin signaling could aggravate Alzheimer’s disease (AD)^13,21–26^. Therefore, it is important to understand the molecular mechanisms by which Reelin is regulated. Of particular importance is understanding how Reelin signaling is turned off because inhibition of such a mechanism would strengthen Reelin signaling and may help improve brain disorders caused by diminished Reelin function.

Secreted signaling proteins can be inactivated or disabled by a variety of mechanisms, including internalization by its receptor, sequestration by an inactive (decoy) binding partner, and degradation. Internalization via ApoER2 and VLDLR^27,28^ and a dominant-negative effect of the secreted extracellular domain of ApoER2^29^ play a role in limiting Reelin signaling. Reelin protein (approximately 430 kDa as a monomer) contains three specific cleavage sites (N-t, C-t, and WC; reviewed in ^7^). N-t cleavage occurs between Pro1244 and Ala1245 to generate two fragments with molecular masses of 160 kDa (NR2) and 270 kDa (Fig. 1A)^30,31^. C-t cleavage occurs between Ala2688 and Asp2689, resulting in two fragments with molecular masses of 330 kDa (NR6) and 100 kDa (Fig. 1A)^32^. WC cleavage occurs between Arg3455 and Ser3456, which is only six amino acid residues from the C-terminus and does not affect the apparent mobility in SDS-PAGE^14^. N-t cleavage dramatically decreases Dab1 phosphorylation by Reelin^30,31^, while C-t or WC cleavage modulate it^14,32^.

**Figure 1.**
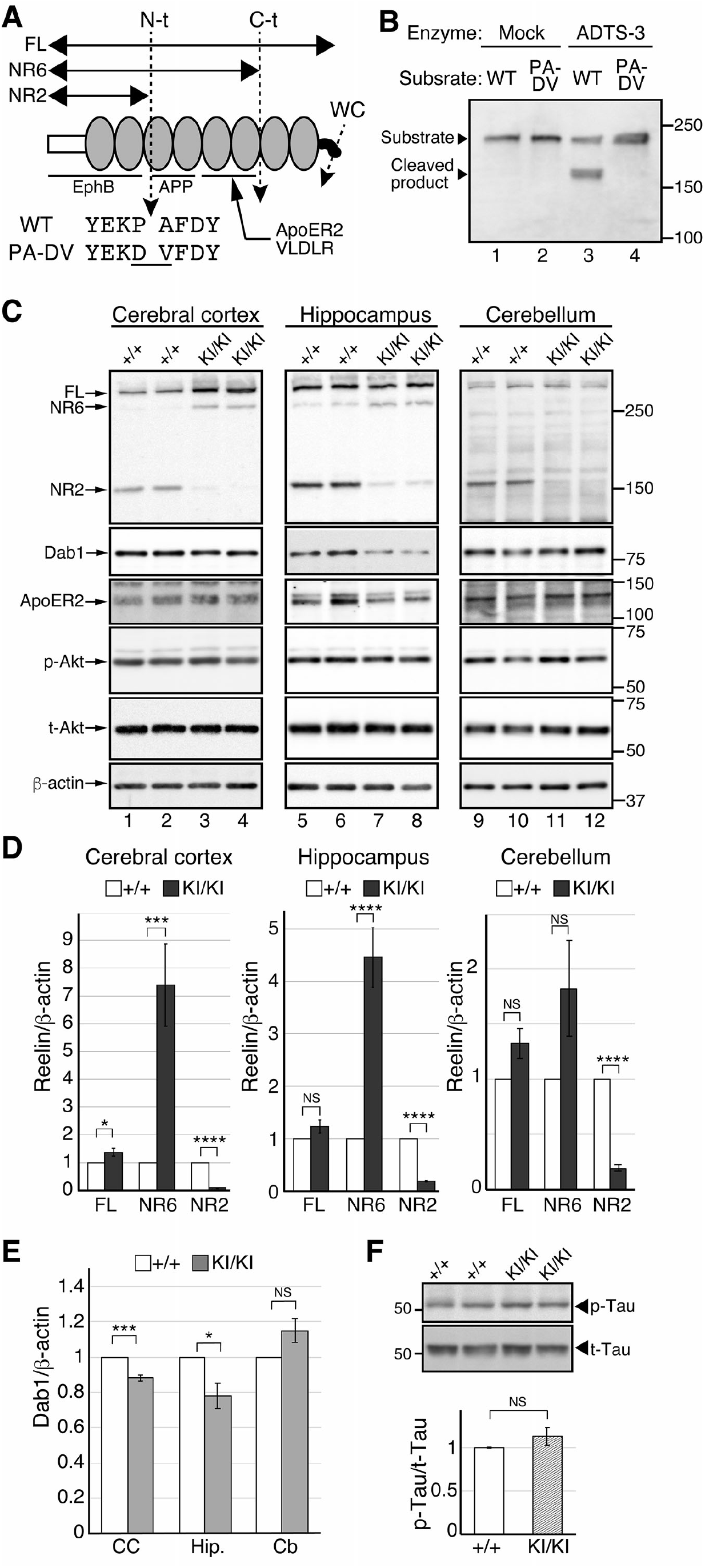
N-t cleavage of Reelin markedly decreased in the embryonic brain of the PA-DV KI mice. **(A)** Schematic drawing of the full-length Reelin protein (FL), its cleavage sites (N-t, C-t, and WC), and its cleaved products (NR6, NR2). The white box and black curve indicate N- and C-terminal regions, respectively. The gray ovals indicate Reelin repeats. Binding regions of known Reelin receptors are indicated with lines. The cleavage sites are indicated by dotted lines with arrowheads. The amino acid sequences around the N-t cleavage site of wild-type (WT) and PA-DV KI mice (PA-DV) are shown below the arrow indicating the N-t cleavage site. The changed residues are underlined. WC, cleavage within the C-terminal region. **(B)** ADAMTS-3 did not cleave the PA-DV sequence. An artificial substrate NR3-MycHis^12,60^ (lanes 1 and 3) or its PA-DV mutant (lanes 2 and 4) were incubated with the culture supernatant from HEK293T cells transfected with control vector (Mock, lanes 1 and 2) or the ADAMTS-3 expression vector (ADTS-3, lanes 3 and 4) for 16 h at 37 °C. The reaction mixtures were separated by SDS-PAGE and analyzed by western blotting with anti-Reelin G10. The positions of the NR3-MycHis (substrate) and its cleaved product are indicated by arrowheads. The positions of the molecular mass markers (kDa) are shown on the right side of the panels. **(C)** Western blotting analysis of the indicated regions of the brain of WT (+/+) or PA-DV homozygous (KI/KI) mice at E18.5 using anti-Reelin G10 (top), anti-Dab1 (middle), and anti-β-actin (bottom) antibodies. Each lane represents a sample from an independent mouse of the indicated genotype. The position of each protein is indicated by arrows. Each lane represents a sample from an independent mouse of the indicated genotype. The positions of the molecular mass markers (kDa) are shown on the right side of the panels. p-Akt, phosphorylated Akt; t-Akt, total Akt. **(D)** Quantification of Reelin FL, NR6, and NR2. White and dark-gray bars indicate the amount of each protein in the +/+ and KI/KI mice, respectively. **(E)** Quantification of Dab1. The white and light-gray bars indicate the amount of Dab1 protein in the +/+ and KI/KI mice, respectively. CC, cerebral cortex; Hip., hippocampus; Cb, cerebellum. **(F)** Western blotting analysis of Tau phosphorylation. The lysates of the cerebral cortex from the indicated mice were separated by SDS-PAGE and analyzed by western blotting with anti-phosphorylated Tau (AT8, top panel) or anti-Tau (Tau5, bottom panel) antibodies. The positions of the molecular mass markers (kDa) are shown on the left side of the panels. p-Tau: phosphorylated Tau, t-Tau: total Tau. The data are shown as the mean ± SEM and were analyzed by two-tailed Student’s t-test. n = 8. \**p* < 0.05, \*\*\**p* < 0.001, and \*\*\*\**p* < 0.0001. NS, not significant (*p* ≥ 0.05).

We recently identified ADAMTS-3 (<u>a</u> <u>d</u>isintegrin <u>a</u>nd <u>m</u>etalloproteinase with <u>t</u>hrombo<u>s</u>pondin motifs 3) as the major protease that mediates N-t cleavage in the embryonic and early postnatal cerebral cortex and hippocampus^12^. More importantly, the results from ADAMTS-3 knock-out (KO) mice indicated that N-t cleavage is the primary mechanism of Reelin inactivation in these tissues^12^. However, ADAMTS-3 was not the only protease to inactivate Reelin; its cleavage still occurred in the cerebral cortex and hippocampus of ADAMTS-3 KO mice^12^. The neuronal dendrites contained more branches and were elongated in the cerebral cortex of ADAMTS-3 conditional KO mice compared to the control mice^12^. We speculated that this effect was due to decreased Reelin cleavage, but it remained possible that there was a critical ADAMTS-3 substrate(s) that somehow affected dendrite morphology. To exclude this possibility, it was necessary to inhibit Reelin cleavage by an alternative approach that did not involve knocking out or inhibiting ADAMTS-3 or other proteases. In fact, we recently found that ADAMTS-2 contributes to Reelin cleavage^33^. Because ADAMTS-2 has many substrates and its KO mice show some Reelin-independent phenotypes in non-neuronal tissues^33,34^, even if ADAMTS-2/3 double KO mice were to show any brain phenotype, it would be impossible to conclude that it is caused by the decrease of Reelin cleavage.

In the present study, we generated Reelin knock-in (KI) mice in which the amino acid residues for the N-t cleavage site of Reelin were substituted by “uncleavable” ones. Characterization of these mice confirmed that N-t cleavage regulates dendritic branching and elongation in the cerebral cortex, an observation obtained using ADAMTS-3 conditional KO mice^12^. More importantly, it also revealed an unexpected role of N-t cleavage in hippocampal development.

## Results

### Establishment of cleavage-resistant Reelin knock-in mice

We attempted to introduce two base changes that would convert Pro1244 to Asp in Reelin using the CRISPR-Cas9 system with a single-stranded DNA fragment (see Methods). We obtained 39 phenotypically normal mice and sequenced the targeted genomic region. The mutation efficiency was quite low (2.6%; 38 of 39 mice had no intended mutation). The remaining mouse carried not only the intended mutations but also an additional mutation at the cleavage site (CCA GCT to <u>GA</u>C G<u>T</u>T; underlined, intended; double-underlined, unintended) that converted Pro1244-Ala1245 residues to Asp-Val (Fig. 1A). This mouse was referred to as PA-DV KI mouse. We confirmed that this conversion makes Reelin protein completely uncleavable by recombinant ADAMTS-3 (Fig. 1B). There were no obvious abnormalities in the growth, fertility, or apparent behavior of PA-DV KI mice compared to the control mice.

Our previous work demonstrated that ADAMTS-3 is the primary enzyme that mediates N-t cleavage of Reelin in the cerebral cortex and hippocampus of the embryonic and early postnatal mouse^12^. However, other enzyme(s) mediating N-t cleavage and the physiological significance of this process after these developmental stages remained unknown. In the current study, we investigated whether N-t cleavage of Reelin decreased in the brain of PA-DV KI mice at embryonic day 18.5 (E18.5), at postnatal day 7 (P7), and 2-month-old.

At E18.5, the amount of NR2 in the cerebral cortex of homozygous PA-DV KI mice (KI/KI) decreased more than 90%, while those of full-length Reelin (Reelin-FL) and NR6 increased compared to those of wild-type (WT) littermates (+/+; Fig 1C, lanes 1-4, and Fig. 1D, left graph). The amount of Dab1 slightly but significantly decreased in the cerebral cortex of the embryonic KI/KI mice (Fig 1C, lanes 1-4, and Fig. 1E, CC). In the hippocampus of the embryonic KI/KI mice, the amount of NR2 decreased, that of NR6 increased, and that of Reelin-FL did not change (Fig 1C, lanes 5-8, and Fig. 1D, middle graph). The amount of Dab1 slightly but significantly decreased (Fig 1C, lanes 5-8, and Fig. 1E, Hip.). These results indicated that N-t cleavage of Reelin was severely abrogated in the cerebral cortex and hippocampus of the embryonic KI/KI mice. They also suggested that the activity of Reelin was augmented in these tissues of KI/KI mice. In the cerebellum of the embryonic KI/KI mice, the amount of NR2 decreased while those of Reelin-FL and NR6 did not significantly change (Fig 1C, lanes 9-12, and Fig. 1D, right graph). Importantly, the amount of Dab1 did not change, too (Fig 1C, lanes 9-12, and Fig. 1E, Cb). Therefore, although N-t cleavage decreased in the cerebellum of the embryonic KI/KI mice, the activity of Reelin to induce Dab1 phosphorylation was not affected in the embryonic cerebellum. The amount of ApoER2 and phosphorylated Akt in the cerebral cortex, the hippocampus, and the cerebellum of KI/KI mice were at the same level as those in WT littermates (Fig. 1C). We previously reported that ADAMTS-3 deficiency decreases Tau phosphorylation in the embryonic cerebral cortex^12^. However, the phosphorylation level of Tau in the cerebral cortex of the E18.5 KI/KI mice was approximately the same as that of WT littermates (Fig. 1F).

### The PA-DV mutation reduces N-t cleavage of Reelin in the postnatal brain

We investigated whether N-t cleavage of Reelin decreased in the brain of PA-DV KI mice at postnatal day 7 (P7). The amounts of NR2 in the cerebral cortex of heterozygous (+/KI) and homozygous (KI/KI) PA-DV KI mice were approximately 40% and 85% lower than that of WT littermates, respectively (Fig. 2A, lanes 1-6, and Fig. 2B, left graph). The amount of Reelin-FL and NR6 slightly increased in the cerebral cortex of KI/KI mice (Fig. 2A, lanes 1-6, and Fig. 2B, left graph). The amount of Dab1 significantly decreased in this brain region of the KI/KI mice compared to that of WT littermates (Fig. 2A, lanes 1-6, and Fig. 2E). The same tendency was observed in the hippocampus (Fig. 2A, lanes 7-12, Fig. 2B, and Fig. 2E). These results showed that Reelin signaling was augmented in the cerebral cortex and hippocampus of PA-DV KI mice at early postnatal stages. In contrast, the amount of Dab1 was little affected in the cerebellum of the PA-DV KI mice (Fig. 2A, lanes 13-18, and Fig. 2E) even though N-t cleavage was inhibited (Fig. 2A, lanes 13-18, and Fig. 2B, right graph). Therefore, the inhibition of N-t cleavage did not seem to augment Reelin signaling in the cerebellum significantly.

**Figure 2.**
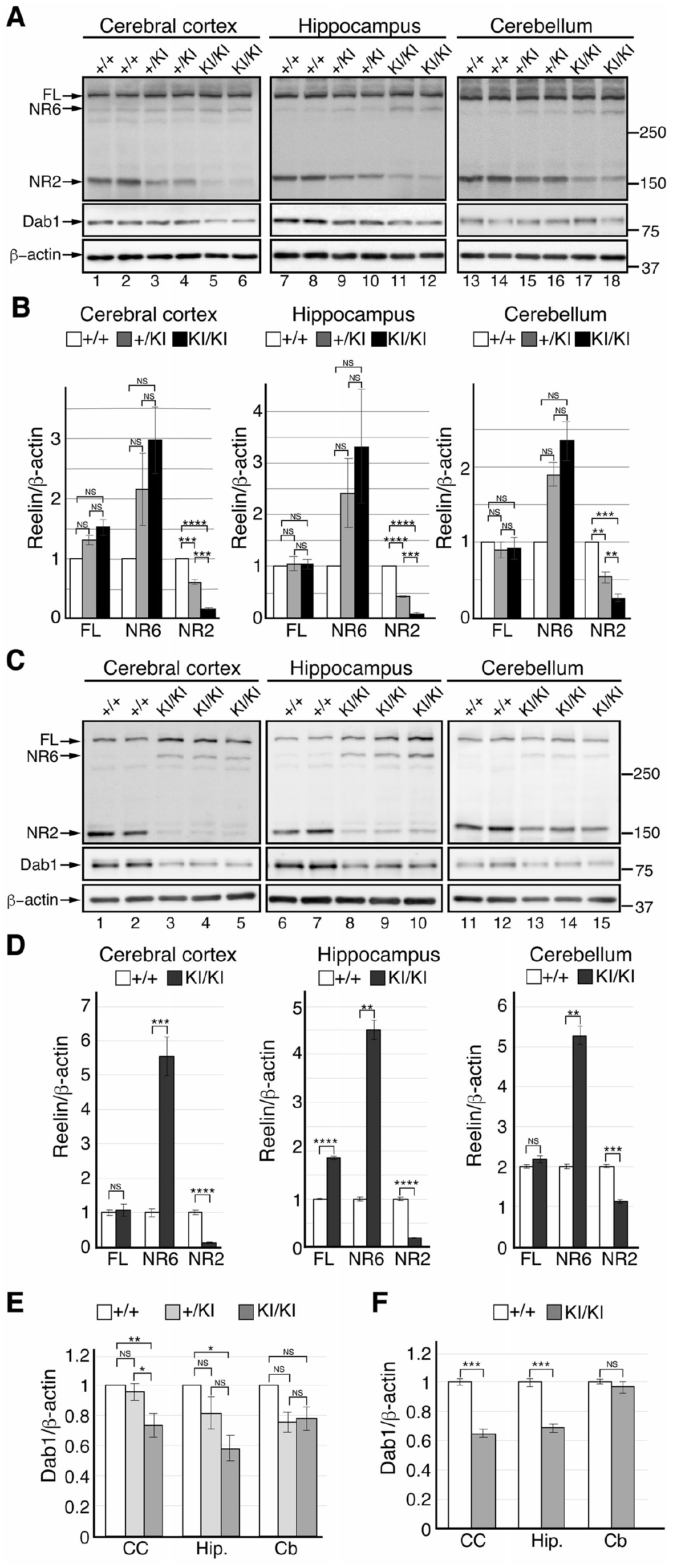
N-t cleavage of Reelin decreased in the postnatal brain of the PA-DV KI mice. (**A, C**) Western blotting analysis of the indicated regions of the brain of WT (+/+), PA-DV heterozygous (+/KI), or homozygous (KI/KI) mice at P7 (**A**) or 2-month-old (**C**) using anti-Reelin G10 (top), anti-Dab1 (middle), and anti-β-actin (bottom) antibodies. Each lane represents a sample from an independent mouse of the indicated genotype. The position of each protein is indicated by the arrow. The positions of the molecular mass markers (kDa) are shown on the right side of the panels. (**B, D**) Quantification of Reelin FL, NR6, and NR2 of the indicated regions of the brain of WT (+/+), PA-DV heterozygous (+/KI), or homozygous (KI/KI) mice at P7 (**B**) or 2-month-old (**D**). White, light-gray, and black bars indicate the amount of each protein in the +/+, +/KI, and KI/KI mice, respectively. *n* = 8. (**E, F**) Quantification of Dab1. of the indicated regions of the brain of WT (+/+), PA-DV heterozygous (+/KI), or homozygous (KI/KI) mice at P7 (**E**) or 2-month-old (**F**). White, light-gray, and black bars indicate the amount of Dab1 protein in the +/+, +/KI, and KI/KI mice, respectively. *n*=8. CC: cerebral cortex, Hip.: hippocampus, Cb: Cerebellum. The data are shown as the mean ± SEM and were analyzed by One-way ANOVA followed by the Tukey-Kramer ad hoc test (**B**, **E**) or two-tailed Student’s t-test (**D**, **F**). n = 6 (**B**, **E**), n = 8 for WT mice and n = 10 for PA-DV homozygous mice (**D**, **F**). \**p* < 0.05, \*\*\**p* < 0.001, and \*\*\*\**p* < 0.0001. NS, not significant (*p* ≥ 0.05).

We next investigated the brain of adult (2-month-old) mice. The overall tendency was the same as observed in the P7 brain. In both the cerebral cortex and hippocampus, N-t cleavage was strongly abrogated (Fig. 2C, lanes 1-10, and Fig. 2D, left and middle graphs) and the amount of Dab1 decreased (Fig. 2C, lanes 1-10, and Fig. 2F) in KI/KI mice. However, N-t cleavage only slightly decreased in the cerebellum (Fig. 2C, lanes 11-15, and Fig. 2D, right graph), and the amount of Dab1 was approximately the same as in WT mice there (Fig. 2C, lanes 11-15, and Fig. 2F). Therefore, N-t cleavage appeared to significantly contribute to the inactivation of Reelin in the adult cerebral cortex and hippocampus, but not in the adult cerebellum.

### The structures of the cerebral cortex and cerebellum of KI/KI mice are mainly normal

We investigated the structures of the brain in KI/KI mice by immunohistochemistry. Localization of Reelin did not differ between WT and KI/KI mice at E18.5 (Fig. 3A and 3B). Immunostaining with anti-Brn1 (a marker of layer II to V, Fig. 3C and 3D), anti-Ctip2 (a marker of layers V and VI, Fig. 3E and 3F), and anti-Tbr1 (a marker of layer VI, Fig. 3E and 3F) antibodies revealed that the neocortical layer formation of KI/KI mice is largely normal at E18.5. At P2, the localization of Reelin did not differ between WT and KI/KI mice (Fig. 3G and 3H). Around this developmental stage, the primitive cortical zone (PCZ) is formed just beneath the Reelin-expressing marginal zone^35^. The formation of the PCZ is dependent on Reelin and Dab1^35^ and disturbed in mice that lack the Reelin C-terminal region^14,15^ or VLDLR^36^. We suspected that the PCZ might be altered in KI/KI mice; however, this region formed normally in KI/KI mice (Fig. 3I and 3J). Localization of Reelin was also similar in WT and KI/KI mice at P7 (Fig. 3K and 3L). Immunostaining with anti-Ctip2 (layer V and VI) revealed that these neurons of the cerebral cortex were positioned normally in KI/KI mice (Fig. 3M and 3N). There were no significant abnormalities in the positions of Cux1+ (layers II to IV, Fig. 3O and 3P) and Tbr1+ (layer VI, Fig. 3Q and 3R) neurons of the cerebral cortex of KI/KI mice. The cerebellar structure in KI/KI mice also appeared normal at both P7 (Fig. 3S and 3T) and 2 months of age (Fig. 3U and 3V). Therefore, N-t cleavage did not appear to play essential roles in the layer formation of the cerebral cortex or cerebellum. Interestingly, we noticed that some of the Ctip2+ neurons of the CA1 region of the hippocampus were located outside of the pyramidal layer (stratum pyramidale, SP) and were present in the molecular layer (stratum oriens, SO) (Fig. 3N and Fig.4A, yellow box). This point was further investigated below.

**Figure 3.**
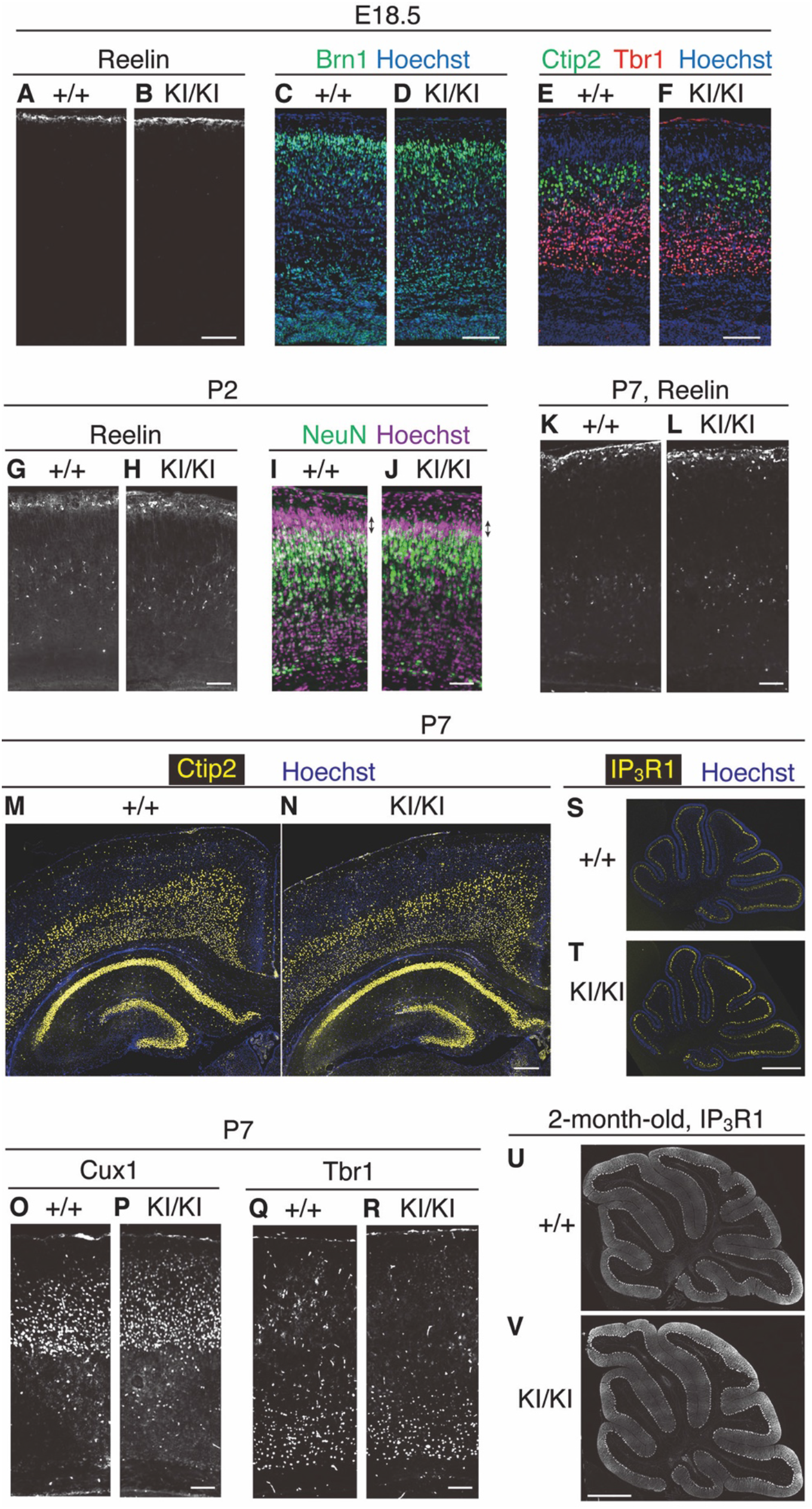
The structures of the cerebral cortex and cerebellum are normal in PA-DV KI mice. (**A**-**F**) Immunohistochemistry of coronal brain sections of the indicated mice at E18.5. Anti-Reelin AF3820 (**A**, **B**), anti-Brn1 (green, **C**, **D**), anti-Ctip2 (green, **E**, **F**), and anti-Tbr1 (red, **E**, **F**) antibodies were used. Nuclei were stained using Hoechst 33342. Scale bar, 100 μm. (**G**-**J**) Immunohistochemistry of coronal brain sections of the indicated mice at P2. Anti-Reelin AF3820 (**G**, **H**) and anti-NeuN (green, **I**, **J**) antibodies were used. Nuclei were stained using Hoechst 33342 (**I**, **J**). Arrows indicate primitive cortical zone (**I**, **J**). Scale bar, 100 μm. (**K**-**R**) Immunohistochemistry of coronal brain sections of the indicated mice at P7. Anti-Reelin AF3820 (**K**, **L**), anti-Ctip2 (yellow, M, N), anti-Cux1 (**O, P**), and anti-Tbr1 (**Q**, **R**) antibodies were used. Nuclei were stained using Hoechst 33342. Scale bars, 100 μm (**K**, **L**, **O-R**), 300 μm (M, N). (**S**-**V**) Immunohistochemical analysis of the sagittal section of the cerebellum of mice at P7 (**S, T**) or 2 months (**U, V**) using anti-inositol-1,4,5-trisphosphate receptor type 1 (IP_3_R1), a marker of Purkinje cells. Nuclei were stained using Hoechst 33342 (blue). Scale bars, 300 μm (**S, T**), 1 mm (**U, V**).

### Some hippocampal neurons were ectopically located in KI/KI mice, which could be rescued by the hemizygous deficiency of the Reelin gene

We observed some ectopic neurons located in the SO, where some Reelin-expressing neurons were also identified (Fig. 4A, yellow box). Quantification of Ctip2+ neuron localization showed that approximately 10% of these neurons were in the SO of KI/KI mice (Fig. 4H, black bars) while such localization was rarely observed in the control mice (Fig. 4H, white bars). This result was unexpected because a similar phenomenon was reported in the mice with attenuated Reelin signaling^14,37,38^ but not in ADAMTS-3 KO mice^12^.

**Figure 4.**
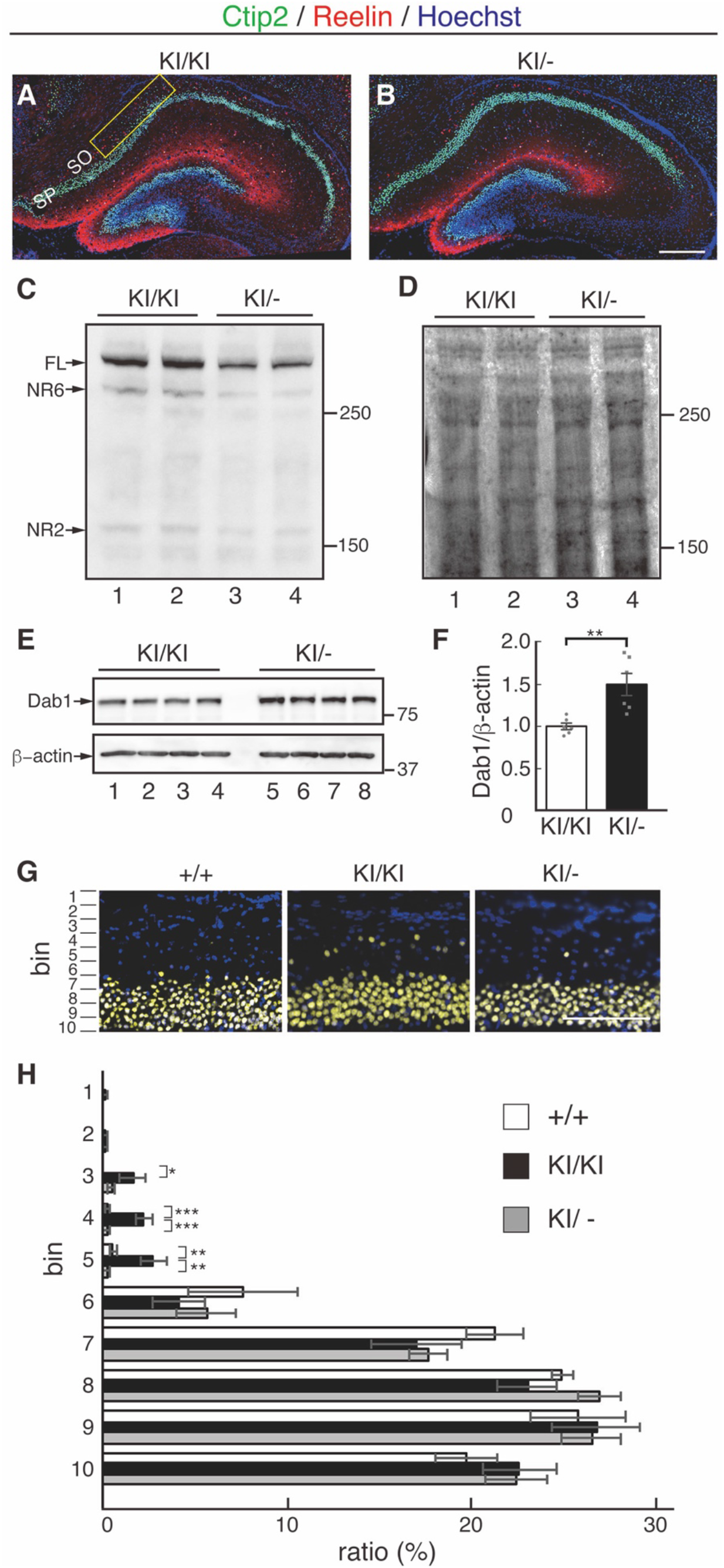
Excess Reelin signaling in the hippocampus causes ectopic neuronal localization in PA-DV KI mice. (**A, B**) Coronal sections of the hippocampus at P7 were immunostained with anti-Ctip2 (green) and anti-Reelin AF3820 (red) antibodies. Nuclei were stained using Hoechst 33342 (blue). Ectopic neurons were observed in the hippocampus of KI/KI mice (**A**, yellow box) but much less in that of KI/− mice (**B**). Scale bar, 300 μm. (**C, D**) Western blotting of hippocampus lysates from KI/KI and KI/− mice at P7 using anti-Reelin antibody G10 (**C**). Ponceau S staining showed approximately equal amounts of protein in each lane (**D**). The positions of full-length Reelin (FL) and the cleaved fragments (NR6, NR2) are indicated by the arrows in **C**. The positions of the molecular mass markers (kDa) are shown on the right. (**E, F**) Western blotting of hippocampus lysates from four independent KI/KI and KI/− mice at P7 using anti-Dab1 (top) and anti-actin (bottom) antibodies (**E**). The positions of the molecular mass markers (kDa) are shown on the right. Quantification of the blots is presented in **F**. (**G, H**) Quantification of the position of Ctip2+ cells in the hippocampus. Scale bar, 100 μm. Data are presented as the mean± SEM and analyzed by two-tailed Student’s t-test (**F**), or One-way ANOVA followed by the Tukey-Kramer ad hoc test (**G**). n = 6. *p < 0.05, **p < 0.01, ***p < 0.001.

Two possible scenarios could explain this phenotype. First, excess Reelin signaling caused the ectopic localization of the Ctip2+ neurons. Alternatively, the cleaved fragment (e.g., NR2) was involved in the positioning of Ctip2+ neurons, and its decrease led to the observed abnormality. To discern between these scenarios, we intercrossed KI/KI mice with Reelin heterozygous deficient (+/−) mice to generate KI/− mice. As expected, the levels of Reelin and its fragments in the hippocampus of KI/− mice were approximately half that of KI/KI mice (Fig. 4C and 4D). Importantly, the amount of Dab1 in the hippocampus of KI/− mice markedly increased compared to that of KI/KI mice (Fig. 4E and 4F), indicating that Reelin signaling in KI/− mice decreased compared to KI/KI mice. Immunohistochemistry showed that almost all of the Ctip2+ neurons were in the correct position in the hippocampus of KI/− mice (Fig. 4B, 4G, and 4H, gray bars). Therefore, these data strongly suggested that excess Reelin signaling, but not a decrease in the amount of the Reelin fragment, caused the ectopic localization of neurons in the hippocampus.

### The dendrites of PA-DV KI mice are more branched and elongated

Reelin is known to induce elongation and branching of neuronal dendrites^39,40^. Indeed, both the total length and the number of branches increased in the layer V neurons of the cerebral cortex of ADAMTS-3 conditional KO mice^12^. However, the phenotypes of ADAMTS-3 conditional KO mice might have been due to other important substrates and not to increased Reelin signaling. We thus investigated the neuronal morphology of the layer V neurons of the cerebral cortex of KI/KI mice at P14 using Golgi-Cox staining (Fig. 5A). We found that both the branching (Fig. 5B) and growth (Fig. 5C) of secondary dendrites of the layer V neurons were augmented in KI/KI mice. Therefore, the upregulation of Reelin signaling by the inhibition of N-t cleavage was sufficient to increase dendritic growth and branching in the postnatal cerebral cortex.

**Figure 5.**
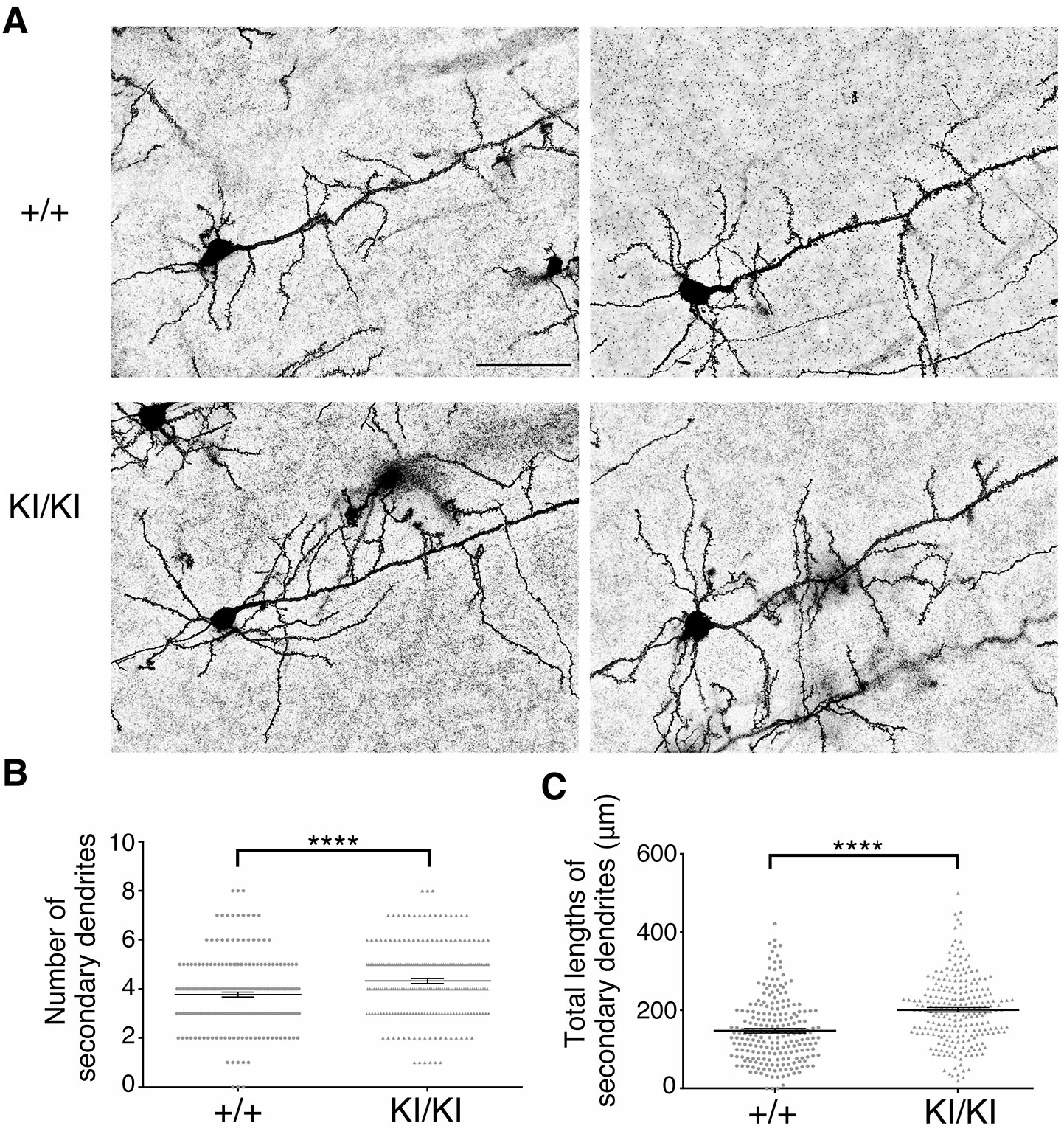
The dendrites of layer V neurons in PA-DV KI mice showed increased branching and elongation. **(A)** Representative images of Golgi-stained layer V neurons in the somatosensory cortex of control (+/+) and KI/KI mice at P14. Scale bar, 50 μm. (**B**) The number of secondary dendrites that were branched from the primary dendrite within 80 μm of the soma. (**C**) The total length of the secondary dendrites. For both **B** and **C**, the cell numbers analyzed were 219 and 230 for +/+ and KI/KI mice, respectively, from three independent mice. The data are shown as the mean ± SEM and analyzed by two-tailed Student’s t-test. ****p < 0.0001.

## Discussion

This study produced several novel findings concerning the regulation and function of Reelin signaling in the brain. First, N-t cleavage indeed inactivated Reelin *in vivo*. Second, the contribution of proteolytic cleavage to the regulation of Reelin function is different between the cerebral cortex/hippocampus and the cerebellum, because the amount of Dab1 did not change in the cerebellum of KI/KI mice. Third, the modulation of Reelin signaling by N-t cleavage is necessary for the proper formation of the hippocampus. Fourth, the phenotype of dendrites previously reported in ADAMTS-3 KO mice was replicated in KI/KI mice, strongly suggesting that the inhibition of Reelin cleavage, not other unidentified substrates, was responsible for the effects of ADAMTS-3 deficiency on the neuronal dendrites. Taken together, these results demonstrated that specific proteolytic cleavage is an integral regulatory mechanism of Reelin function.

The regulation of Reelin activity by proteolytic cleavage has been studied by several groups on several different systems. Goffinet and colleagues performed pioneering works, finding that Reelin is processed by metalloproteases^41^ and the cleavage sites lie roughly between Reelin Repeat 3 and 6^42^. They also showed that the application of a broad-spectrum metalloproteinase inhibitor abrogated the neuronal migration *in vitro*^43^. Although cerebral cortical neuronal migration appears normal in ADAMTS-3 KO mice^12^ and in PA-DV KI mice, it is still possible that a fragment of Reelin plays a role in certain circumstances, because N-t cleavage is not fully abolished in these mice. The importance of N-t cleavage as the liberation of the “active Reelin fragment” was also proposed by Tinnes and colleagues in the maintenance of dentate granule cell positioning after hyperexcitation induced by kainic acid^44^. It would be interesting to see if this phenomenon is reduced in PA-DV KI mice.

The difference in the amount of NR2 between the control and KI/KI mice was striking regardless of the investigated tissues and time points; the change in FL was much smaller (Fig. 1 and Fig. 2). This tendency was also observed in ADAMTS-3 KO mice^12^. There are two reasons for this apparent discrepancy. First, most of Reelin-FL resides in the intracellular pre-secretory compartment, which is not affected by extracellular proteolysis^41^. Second, both Reelin-FL and NR6, but not NR2, can bind to ApoER2 and VLDLR, leading to their internalization and metabolism^45^. Thus, NR2 levels are the best indication of N-t cleavage, and the apparent discrepancy in the amounts of Reelin-FL and NR6 is not important in the scope of the present study.

The amount of Dab1 of the cerebral cortex and hippocampus was lower in KI/KI mice than in control mice at all the investigated time points (Fig. 1E and 2E). These results indicate that Reelin activity is augmented in these tissues. On the other hand, it was quite unexpected that the amount of Dab1 was unaffected in the cerebellum of KI/KI mice (Fig. 1E and 2E). Reelin and Dab1 are expressed in the granular neurons and Purkinje cells (PCs) of the cerebellum, respectively^46^. To the best of our knowledge, whether Reelin can induce phosphorylation and degradation of Dab1 in PCs has never been rigorously investigated. In the cerebellum of mice that lack the C-terminal region of Reelin, some PCs are ectopically located and contain a higher amount of Dab1 protein^47^. This observation provides indirect evidence that Reelin-induced Dab1 degradation does occur in PCs and raises the question as to what could make a difference between the cerebellum and cerebral cortex or hippocampus. It is possible that some other signaling molecules or pathways control the amount of Dab1 in the cerebellum. Dab1 is downstream of other signaling molecules, such as APP^48^, LRP1^49^, and DCC^50^. These molecules may affect the amount of Dab1 in the postnatal cerebellum. Another possibility is that the cleaved fragment of Reelin, presumably the central fragment^42^, retains full activity in the cerebellum, unlike in the cerebral cortex or hippocampus. It was recently reported that the unstable intronic ATTTC repeat mutation in the human DAB1 gene leads to overexpression of Dab1 protein, resulting in spinocerebellar ataxia type 37^51^. Interestingly, the structural abnormality is found only in the cerebellum in these patients, and they show only cerebellar dysfunctions^51^. It is thus likely that excess Reelin-Dab1 signaling damages the cerebellum only, which in turn suggests the existence of a mechanism that strongly shuts down Reelin activity in the healthy cerebellum. Finally, while N-t cleavage of Reelin and the amount of Dab1 decreased in the cerebral cortex and hippocampus of postnatal ADAMTS-2 KO mice, exactly the opposite results were obtained in the cerebellum^33^. All of these observations indicated that the mechanism of N-t cleavage as well as the impact of Reelin on the amount of Dab1in the cerebellum are different from those in the cerebral cortex or hippocampus.

We also noticed that the amount of NR2 in the cerebral cortex was much lower in KI/KI mice than in ADAMTS-3 KO mice^12^. Therefore, N-t cleavage was much more severely inhibited in KI/KI mice than in ADAMTS-3 KO mice. Nonetheless, the difference of the Dab1 amount between KI/KI mice and controls was less striking than in the case of ADAMTS-3 KO mice^12^. This may explain why the level of Tau phosphorylation in the embryonic cerebral cortex, which was markedly down-regulated in ADAMTS-3 KO mice^12^, was not significantly affected in KI/KI mice (Fig. 1E). Alternatively, ADAMTS-3 deficiency may have a Reelin-independent function that affects the phosphorylation of Dab1 and Tau in the embryonic stages.

In the present study, we showed that some of the pyramidal neurons are mislocalized in the SO of the hippocampus of these mice. This result was unexpected because a similar phenomenon was reported in mice with attenuated Reelin signaling^14,37,38^ but not in ADAMTS-3 KO mice^12^ or mice overexpressing Reelin in adult excitatory neurons^52^. We first hypothesized that the decrease in the levels of the Reelin fragments might be responsible for this abnormality, which had been proposed for cerebral cortical development^42,43^ and the maintenance of dentate gyrus structure^17,44^. However, our results showed that this was not the case. The mislocalization was caused by excess Reelin signaling. Compared to ADAMTS-3 KO mice^12^, the decrease in the NR2 fragment level was more prominent in KI/KI mice, strongly suggesting that Reelin signaling is much more augmented in KI/KI mice than in ADAMTS-3 KO mice. This result likely explains why KI/KI mice, but not ADAMTS-3 KO mice, had a structural abnormality in the hippocampus. To the best of our knowledge, this is the first example of a negative effect of excess Reelin signaling in the developing brain. More importantly, it revealed that N-t cleavage is physiologically relevant for layer formation in the hippocampus. It is also worthwhile to mention the recent report by Dlugosz and colleagues that the central fragment of Reelin may have its specific (and presumably Dab1-independent) role^53^. This mechanism may be involved in the formation of the hippocampus.

Reelin has been implicated in the onset or aggravation of many neuropsychiatric and neurodegenerative disorders^4,6–9,16^. In almost all of the cases, the Reelin hypoactivity worsens them^1,16^. Therefore, increasing Reelin signaling may be an approach to prevent or improve these disorders. In this study, we showed that inhibition of N-t cleavage could solely augment Reelin signaling and induce branching and elongation of neuronal dendrites in the cerebral cortex. An inhibitor against proteases mediating this reaction thus could improve the symptoms of disorders in which Reelin hypoactivity is a contributing factor.

Reelin is known to positively regulate neurogenesis^3,52^, synapse formation^1,7,52,54^, and synaptic plasticity^1,7,52,54^. Whether these events are altered in KI/KI mice is currently being investigated. Mice with attenuated Reelin signaling often exhibit behavioral abnormalities^38,55^. It would be interesting to test if the PA-DV mutations could correct or improve these abnormalities. The PA-DV KI mouse provides a new path for investigating increased Reelin function, particularly in the field of neuropsychiatric disorders.

## Methods

### Reagents and antibodies

The following primary antibodies were purchased: anti-Reelin (G10, MAB5364) and anti-NeuN (MAB377) from Millipore; anti-Reelin (AF3820) from R&D Systems; anti-Cux1 (sc-13024) from Santa Cruz Biotechnology; anti-β-actin (AC-15) from Sigma; anti-Ctip2 (ab18465) and anti-Tbr1 (ab31940) from Abcam. Rabbit anti-Dab1 antiserum was made and affinity purified as described previously^27^. Alexa Fluor 488- or Alexa Fluor 594-conjugated secondary antibodies were purchased from Thermo Fisher Scientific.

### DNA Constructs

The expression vector for the artificial substrate NR3-MycHis was described previously^31^. The expression vector for NR3-PA-DV was generated by introducing mutations into the N-t cleavage site of NR3-MycHis.

### Animals

All experimental protocols were approved by the Animal Care and Use Committee of Nagoya City University and performed according to the guidelines of the National Institutes of Health of Japan. Jcl:ICR and C57BL/6N mice were obtained from Japan SLC. Reelin-deficient *reeler* mice (B6C3Fe-a/a-rl) were purchased from Jackson Laboratories and back-crossed with Jcl:ICR mice.

A knock-in mouse strain in which Pro1,244 and Ala1,245 of Reelin were substituted with Asp and Val residues (PA-DV KI mice) was generated as follows: Mutations were introduced into the mouse Reelin gene using the CRISPR/Cas9 system. A linker corresponding to the sgRNA sequence was generated using primers CACCGTTCATAGGGTAATCGAAAGC and AAACGCTTTCGATTACCCTATGAAC and inserted into the Bbs1 site of the px330 plasmid^56^.

This plasmid and synthetic single-strand DNA (GTTATCCAACCGTCTTCCTGTTTCGTTTGTGTTGTCCCTAATAGAACTTCTATGAGAAGGACGCTTTCGATTACCCTATGAACCAAATGAGTGTGTGGCTAATGTTGGCCAATGAAGGC) were injected into the fertilized egg pronucleus of C57BL/6J mice. Genomic DNA was prepared as described previously^14^ and genotype was determined by PCR (33 cycles at 94°C for 30 s, 64°C for 30 s, and 72°C for 30 s) using the following primers: for WT allele, PDKI-WT-21 (CCCTAATAGAACTTCTATGAGAAGCCA) and PDKI-Rev-24 (CCTTACCAATAGAGCTGAAAGGCTTC); for PA-DV KI allele, PDKI-WT-21 and PADV-KI-93 (CCCTAATAGAACTTCTATGA GAAGGACGT). The sizes of the PCR products were 347 bp and 349 bp for WT and PA-DV KI alleles, respectively.

The PA-DV KI mice were back-crossed with either C57BL/6N or with Jcl:ICR at least 8 times. All experiments shown were performed with C57BL/6N background, except for Fig. 4. In Fig. 4, the mice with Jcl:ICR mice were analyzed.

### Immunohistochemistry

Immunohistochemistry was performed as described previously^12,14^. All data shown in the Figures are representative images of at least three independent mice, otherwise indicated.

### Golgi–Cox staining

Golgi–Cox staining was performed as described previously^12^. Briefly, dissected P14 brains were immersed in the Golgi–Cox solution for three days, followed by seven days in PBS containing 30% sucrose. Brains were cut at a thickness of 160 μm sections using a VT1000S vibratome (Leica Microsystems). Sections were incubated in 15% aqueous ammonium hydroxide for 30 min under a fume hood in the dark followed by 30 min in Kodafix solution (CosmoBio). Sections were rinsed with distilled water twice, and dehydrated using an ethanol series and lemosol and mounted with Softmount (Sakura). Layer V neurons which located in the somatosensory area using a light microscope with a 60× lens, and pictures were taken with the BZ-9000 system (Keyence). The primary and secondary dendrite lengths were measured using ImageJ with the NeuronJ plugin (National Institutes of Health) as described previously^12^. The length of secondary dendrites that branched from the primary dendrite within 80 μm of the soma were counted and measured. Researchers were blinded to the genotypes of the mice in all experiments.

### Cell culture and transfection

Cell culture and transfection was performed as previously described^12,57,58^.

### Western blotting

Western blotting was performed as described previously^12,14^. Dissected brains were sonicated in lysis buffer (20 mM Tris-HCl, pH 7.5, 150 mM NaCl, 5 mM EDTA, 1% Triton X-100, 0.1% H_2_O_2_, and 5 mM Na_3_VO_4_). The blots were analyzed with ImageJ and quantified as described previously^59^.

### Cell culture and transfection

The culturing of human embryonic kidney 293T cells, transfection of plasmid DNA using Lipofectoamine2000, and recovery and preservation of the culture supernatants were performed as described previously^58,59^.

### Western blotting

Dissected cerebral cortices, hippocampi, and cerebellums were homogenized in lysis buffer (20 mM Tris-HCl, pH 7.5, 150 mM NaCl, 5 mM EDTA, 1% Triton X-100, 0.1% H_2_O_2_, and 5 mM Na_3_VO_4_)^57^. Insoluble debris was removed by centrifugation (10 min; 13,000 rpm), and the supernatants were collected. Western blotting was performed as described previously^14^. The blots were analyzed with ImageJ and quantified as described previously^59^.

### Statistical analyses

Data are presented as the mean ± the standard error of the mean (SEM). One-way ANOVA followed by the Tukey-Kramer ad hoc test was used to compare three different groups. Two-tailed Student’s test were used to compare the means of two groups. Statistical analyses were performed with Prism6 (GraphPad Software). Statistical significance was represented as *p < 0.05, **p < 0.01, ***p < 0.001, and ****p < 0.0001.

## Acknowledgements

We thank Drs. Tom Curran (Children’s Research Institute at Children’s Mercy Hospital, MO, USA), Junichi Takagi (Osaka University, Japan), and Katsuhiko Mikoshiba (Brain Science Institute, RIKEN, Japan) for generously providing reagents. This work was supported by JSPS KAKENHI (17H03895 and 17K19500 to M.H., 17K08281 to T.K., and 17J10967 to H.Ogino), ACT-M (16im0210602h0001 and 17im0210602h0002) of the Japan Agency for Medical Research and Development (to M.H.), and Ono Medical Research Foundation (to M.H.). H. Ogino is Research Fellow of Japan Society for the Promotion of Science (DC1).

## Author Contributions

E.O., H.Ogino., T.K., and M.H. conceived and designed research; E.O., H.Oishi., T.K, and M.H. established the PA-DV KI mice.; E.O., H.Ogino., T.S., and Y.Y. performed biochemical and molecular biological experiments; E.O., H.Ogino., T.S., Y.Y., and H.T. performed immunohistochemical experiments; E.O., H.Ogino., T.S., Y.Y., and M.H. performed statistical analyses; E.O., H.Ogino., T. K, and M.H. wrote the manuscript.

